# Understanding The Myelin *g* ratio From First Principles, Its Derivation, Uses And Artifacts

**DOI:** 10.1101/2024.09.08.611849

**Authors:** Alexander Gow

**Author notes:** Correspondence: Dr Alexander Gow, Center for Molecular Medicine and Genetics, 3216 Scott Hall, 540 E Canfield Ave, Wayne State University School of Medicine, Detroit, MI, 48201.

## Abstract

In light of the increasing importance for measuring myelin *g* ratios – the ratio of axon-to-fiber (axon + myelin) diameters in myelin internodes – to understand normal physiology, disease states, repair mechanisms and myelin plasticity, there is urgent need to minimize processing and statistical artifacts in current methodologies. Unfortunately, many contemporary studies fall prey to a variety of artifacts, which reduce study outcome robustness and slow development of novel therapeutics. Underlying causes stem from a lack of understanding of the myelin *g* ratio, which has persisted more than a century. An extended exploratory data analysis from first principles (the axon-fiber diameter relation) is presented herein and has major consequences for interpreting published *g* ratio studies. Indeed, a model of the myelin internode naturally emerges because of (1) the strong positive correlation between axon and fiber diameters and (2) the demonstration that the relation between these variables is one of direct proportionality. From this model, a robust framework for data analysis, interpretation and understanding allows specific predictions about myelin internode structure under normal physiological conditions. Further, the model establishes that a regression fit to *g* ratio plots has zero slope, and it identifies the underlying causes of several data processing artifacts that can be mitigated by plotting *g* ratios against fiber diameter (not axon diameter). Hypothesis testing can then be used for extending the model and evaluating myelin internodal properties under pathophysiological conditions (accompanying article). For without a statistical model as anchor, hypothesis testing is aimless like a rudderless ship on the ocean.

## INTRODUCTION

In the decades following the pioneering work on PNS myelinated fiber morphology by Donaldson and Hoke (1905), relationships between Schwann cell-derived compact myelin and ensheathed axons were studied extensively. Relations between conduction velocity (CV), myelin radial thickness, axon diameter, and internodal length area and volume were found to be linear (or close-to-linear) across a broad range of vertebrate species. Common practice was to represent these factors as relations of cross-sectional myelin-axon area ratios or internodal length or axon and fiber diameters (reviewed by Rushton, 1951).

In the mid-1930s, Schmitt and colleagues used X-ray diffraction (Schmitt et al., 1935) and particularly birefringence (Schmitt and Bear, 1937) to delineate myelin fine structure and quantify the axon-myelin relation in the PNS. In a theoretical treatise on the optical properties of myelin, Bear and Schmitt (1936) developed mathematical equations to interpret the birefringence of myelinated axons. Their analysis defined a series of variables with labels in alphabetical order, the last being “*g*” [equation (12)], defined as the ratio of axon and fiber radii, and thereafter equated with the ratio of axon and fiber diameters [equation (18)]. The name of this experimentally measurable variable evolved from “*g*” (Schmitt and Bear, 1937) to “ratio *g*” (Sanders, 1947) to the current myelin “*g* ratio”. And although widely perceived as being correlated with axon (or fiber) diameter, precisely defined *g* ratios from optical and electrical theory (Bear and Schmitt, 1936; Rushton, 1951) or electron microscopy measurements (Schnepp and Schnepp, 1971; Waxman and Bennett, 1972; Price and Sprich, 1975; Waxman and Swadlow, 1976) are essentially constant (i.e. invariant) under normal physiological conditions across a wide range of fiber calibers in the PNS and CNS.

But even a full view of the last 120 years documenting the intimate interrelationship between myelin sheaths and their axons fails to remit the reality that an exhaustive account of the myelin *g* ratio, the most conspicuous means of characterizing myelination, remains incomplete. In particular, the absence of a detailed understanding of the *g* ratio itself and its major implications and applications to normal physiology, has often left the myelin field in a quandary about the processing, analysis and interpretation of myelin internode data. To wit, the field lacks an explicit well-defined model to provide a framework for characterizing physiological and disease-associated changes in myelin or testing pharmacological agents to remedy myelin pathobiology. This is particularly noteworthy in much of the data published in recent decades.

We need to acknowledge that the experimental design and data processing required to generate and interpret *g* ratio plots is deceptively simple. There are multiple incongruities (explored in this study) with experimental measurement errors that have far-reaching implications for data interpretation. For example, the linear correlation between axon and fiber diameters differs on either side of an approximate “threshold” axon diameter. Well below this threshold, myelin is absent and axon diameter is equal to fiber diameter (*g* ratio = 1). Well above the threshold, axons are myelinated and fiber diameter exceeds axon diameter (0 < *g* ratio < 1). In the vicinity of the threshold [ca. 0.2μm in CNS, (Waxman and Bennett, 1972)], similar sized axons may be myelinated or unmyelinated for reasons currently unclear. Thus, the axon-fiber diameter relation is complex. In the current study, such complexity is put aside for subsequent explanation. Hence, experimental data and simulations herein are confined to myelinated fibers.

In this vein, we revisit the original model of the myelin *g* ratio in the novel context of a directly proportional relation between axon and fiber diameters. The model generates a statistical framework that pertains only to myelin internodes under normal physiological conditions, but can be extended to pathological systems using hypothesis testing. Previously published wild type data from adult optic nerve (Dupree et al., 2015) is used to motivate the current study and delineate data processing and interpretation artifacts that are common in the literature. These artifacts are explored using simulations, which clarify interpretations for publishing *g* ratio data from both the PNS and CNS.

## METHODS

### Experimental data and assurances of humane care of mice

Experimental data analyzed in the current project were obtained from Dr Douglas Feinstein, Department of Anesthesiology, University of Illinois at Chicago, and Department of Veterans Affairs, Jesse Brown VA Medical Center (Dupree et al., 2015). The authors stated that all animal procedures were humane and approved by the Institutional Animal Care and Use Committee.

### *g* ratio measurements in adult optic nerve from a control cohort of mice

The axon and fiber diameter data used in this study are from the Sham-treated cohort of mouse optic nerves published by Dupree and colleagues (2015). Briefly, this control cohort was treated with saline emulsified in complete Freund’s adjuvant, including *M. tuberculosis* and pertussis toxin components but not MOG_35-55_ peptide that is used to induce EAE. After seven weeks, the mice were perfused with 4% paraformaldehyde / 5% glutaraldehyde in 0.1M Millonig’s buffer, post fixed with 2% osmium tetroxide and embedded in PolyBed 812 resin. This procedure may cause *g* ratios to be larger than the typical range (i.e. 0.75-0.8) found in the literature. Thin plastic sections of transverse optic nerve were imaged under the electron microscope from 10 randomly selected fields (magnification: 10,000x; camera pixel resolution = 2.27nm) per mouse. Axon and fiber diameter, *g* ratio and myelin “diameter” (equivalent to 2 x the myelin radial thickness) were measured. Unmyelinated axons were not measured.

### Simulations of axon, fiber and myelin diameters and *g* ratios

All simulated values used in this study were generated in Microsoft excel (ver16.66.1). The range of fiber diameters was 0.3-18.8μm with a step size of 0.5μm. Four *g* ratios were generated, {0.64, 0.70, 0.76, 0.82}, with a step size of 0.06 and a grand average *g* ratio = 0.73. These ranges essentially encompass the entirety of fiber diameters and *g* ratios observed in the wild type vertebrate nervous system. From these criteria, axon diameters were computed using the standard *g* ratio equation,

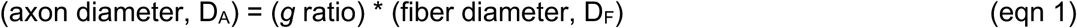

and myelin diameter,

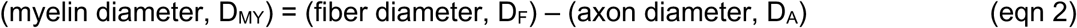

Random sampling of the simulated values was performed using the Excel RAND() function with 0 ≤ *random#* < 1 and a selection threshold > 0.7 (∼30% probability of selection). The selected values were imported into Graphpad Prism v10.3.0 (GraphPad Software, LLC, San Diego, CA) for all subsequent processing.

### Processing of data and simulations for statistics

Microsoft excel was used to collate raw data measurements from TEM micrographs of Sham-treated mice (Dupree et al., 2015), and to compute axon, fiber and myelin diameters, and *g* ratios. These data were imported into Graphpad Prism for all subsequent processing and statistical analyses. Pearson’s correlation coefficient, *r_xy_*, Spearman’s correlation coefficient, *r_S_*, goodness of fit, R^2^, and adjusted P values were computed for unweighted regression fits using Deming linear and nonlinear regression algorithms in Prism. When comparing linear versus polynomial or logarithmic fits to plots of myelin diameter as a relation of fiber or axon diameter, Akaike’s Information Criterion (AICc) was used (Dziak et al., 2020). This analysis is robust against the effects of pseudo-replicate axon and fiber diameter data measured experimentally from all electron micrographs of white matter tracts. Alternatively, the impact of pseudo-replicates on statistical tests has been reduced using data binning or averaging e.g. computing the average *g* ratio for individual mice prior to t-tests.

### Box-Cox Lambda Transformation Series for statistics

This series is a useful approach to determine if nonlinear axon diameter (*y*-axis) transformations [e.g. (*y*-axis)^2^, ln(*y*-axis), 1/(*y*-axis)] might improve the Pearson’s correlation coefficient compared with the value for the untransformed data [i.e. the (*y*-axis)^1^ transformation]. The curve generating the highest Pearson’s coefficient is the best regression curve fit and, most likely, the true relation between the two variables. The transformed data are also plotted against untransformed fiber diameters (*x*-axis) and fit with simple linear regression curves for visual conformation of the best linear correlation. This goal of this procedure can be formulated as follows: if a nonlinear relation exists between the variables, then Pearson’s coefficient will be maximized by the *y*-axis transformation that most likely corresponds with the true relation.

To perform the data transformations, axon diameters of individual mice from the Sham cohort in the EAE study were transformed using the Box-Cox Lambda series (Box and Cox, 1964) according to:

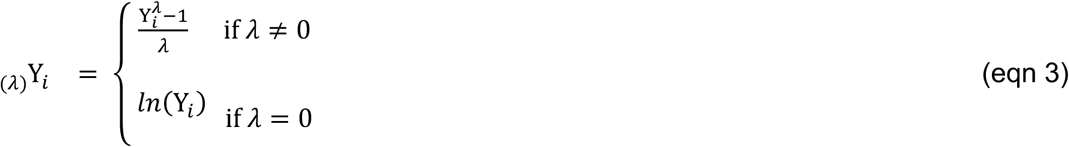

for defined values, λ = {-3, −2, −1, −0.5, 0, 0.5, 1, 2}, where λ = 0 corresponds to ln(Y*_i_*), and _(*λ*)_Y*_i_* are the Box-Cox transformed *y*-axis values. A numerical constant, added to all Y*_i_*, can be convenient if any _(*λ*)_Y*_i_* < 0. Values of _(*λ*)_Y*_i_* are plotted against the corresponding fiber diameter values in a summary plot.

Transformations of *y*-axis data modify the unit of measurement (e.g. from units of μm to μm^2^ if *λ* = 2) and may need to be reverse transformed to recover the original unit of measurement prior to transformation. Such reverse computations are performed according to:

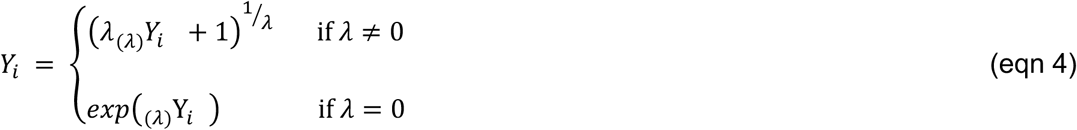

A crude linearity test for the Box-Cox transformation series involves the Spearman’s correlation coefficient, *r_S_*. As a non-parametric test, Spearman’s correlation is largely dependent on the rank-order of the axon-fiber diameter pairs. Because Box-Cox transformations do not change this order, *r_S_* is invariant for all λ values. In this light, Spearman’s coefficient can often be interpreted as a de facto lower limit for linear correlation between two variables. Thus, in conjunction with Pearson’s coefficients, Box-Cox transformations for which *r_xy_* > *r_S_*, often can be interpreted as transformations that improve both the linear association between, and normal distribution of, the variables.

Several statistics – Pearson’s and Spearman’s coefficients, regression curve slopes, *x*- and *y*-intercepts – were extracted from Box-Cox series summary graphs, and plotted against the corresponding λ values to identify the transformation with properties that most similarly match the biology of the axomyelin unit. All Pearson’s correlation coefficients, *r_xy_*, generated in the current study were transformed using Fisher’s Z equation (van Aert, 2023):

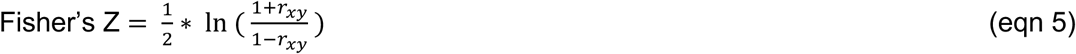

prior to statistical testing using paired t-tests or mixed-effects analysis (similar to repeated measures ANOVA) with Geisser-Greenhouse epsilon correction and Dunnett’s post hoc tests. Regression slopes and *x*- and *y*-intercepts were assessed for consistency with the known biological properties of the axon-fiber diameter plot. The major steps of this analysis are summarized in Table 1.

**Table 1.**
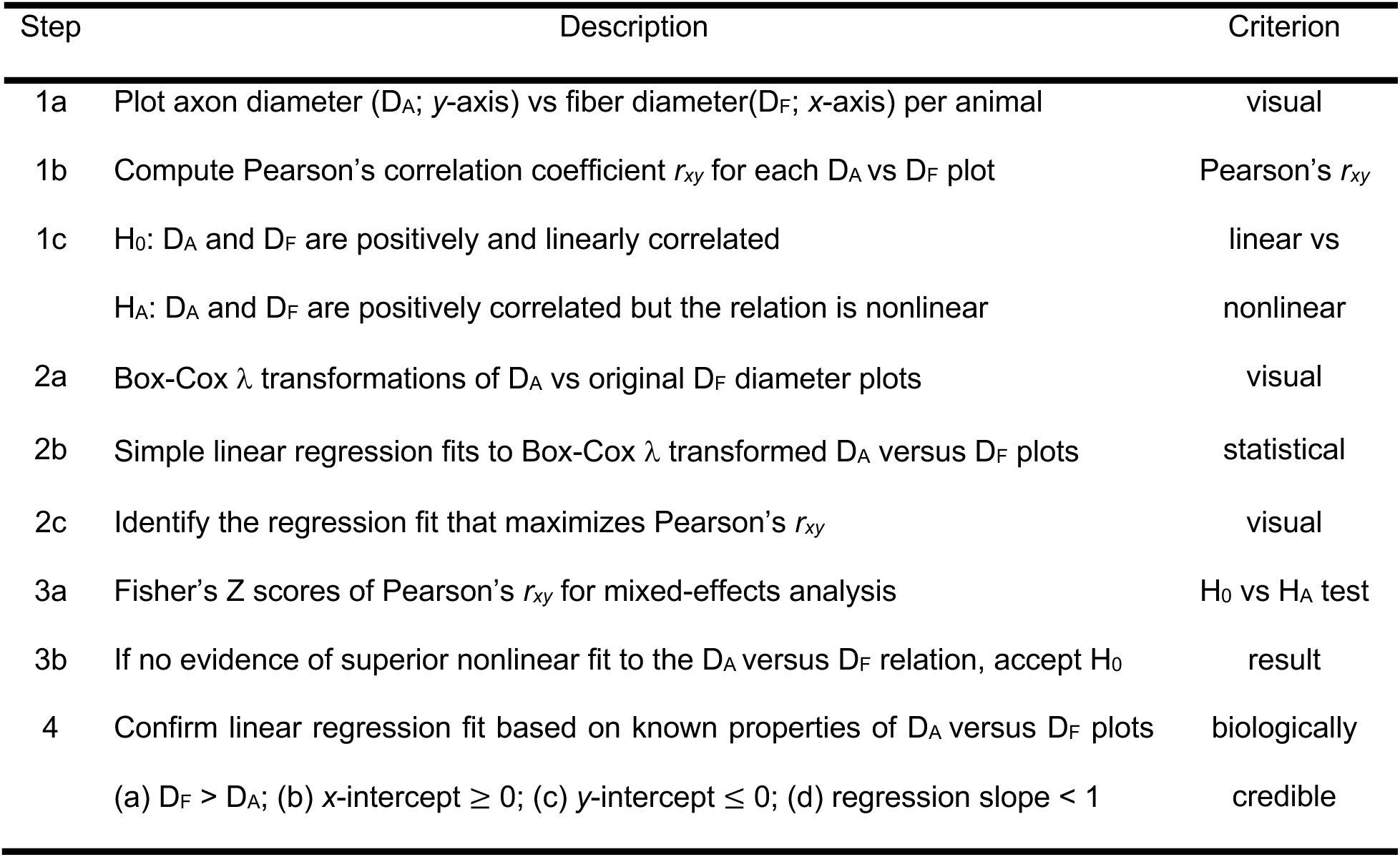
– criteria to determine optimal regression fit to axon-fiber diameter plots.

For the simulations in Fig. 6, linear regression was used for curve fitting, as well as a generalized logarithm model (pink line) according to:

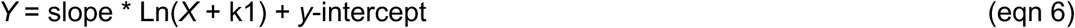

where k1 represents a scaling constant that prevents negative log-transformed values. Outlier detection and removal was not performed.

## RESULTS

The current study uses previously published *g* ratio data from electron micrographs of adult mouse optic nerve (Dupree et al., 2015), summarized using graphs typically seen in the literature. The data are used to motivate the identification and investigation of artifacts in *g* ratio data that generally go unrecognized: unstable simple linear regression slopes and *y*-intercepts, data structure distortions, random sampling effects and the presence of influential data points, which exert undue influence on regression curve fits. Most of these artifacts are associated with or more pronounced when axon diameter is the independent variable for *g* ratio and myelin plots, while corresponding plots using fiber diameter are largely artifact-free.

### Regression fits for *g* ratio plots are unstable

Optic nerve data from three mice in Fig. 1A are plotted with *g* ratio as a relation of fiber diameter. Panel 1B shows the same *g* ratio data plotted against axon diameter, and panel 1C shows the grand average *g* ratio (mean ±S.D.) determined from the median *g* ratios of each mouse. In panels 1A and 1B, the individual mouse data are also fit using linear regression (as is typical in the published literature). Several features are noteworthy.

**Figure 1.**
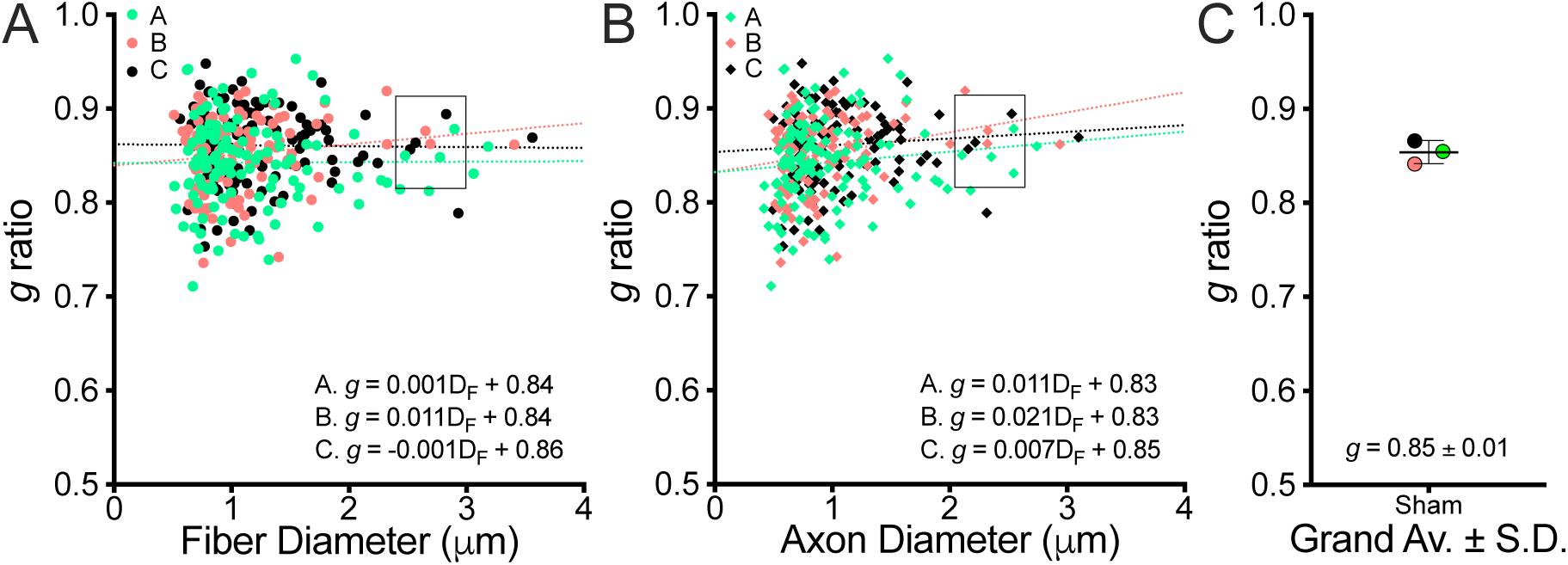
*g* ratios as relations of axon diameter, fiber diameter or grand average from wild type mice Comparisons of **A.** *g* ratio versus fiber diameter plot, **B.** *g* ratio versus axon diameter plot and **C.** grand average (mean ±S.D.) of the Sham mouse cohort from a previously published study (Dupree et al., 2015). The data in (A) and (B) are fit using linear regression for each mouse. Data points in the axon diameter plot are compressed along the *x*-axis relative to the fiber diameter plot (i.e. 0-3.6μm versus 0-3.1) and some data points shift along the *x*-axis relative to other points (compare positions of the green and black data points in the rectangles), while the spread of *g* ratios along the *y*-axis is unchanged. The choice of *x*-axis variable apparently causes the regression slopes to change sign, which may be associated with reordering of the data points along the *x*-axis. Concurrently, *y*-intercepts are unstable.

The most obvious features in panels 1A and 1B are the compression of data points along the *x*-axis in panel 1A relative to 1B (i.e. 0-3.6 versus 0-3.1μm), and the changes in individual regression slopes. The vertical spread of *g* ratios on the *y*-axis is identical in both panels. Surprisingly, the simple substitution of fiber diameter for axon diameter on the *x*-axis affects the regression slopes in several ways. For example, the regression lines are steeper (more positive) in panel 1B than 1A, and the regression slope for mouse C changes sign (negative in 1A, positive in 1B).

Variable regression slopes between mice in published studies are common, and often dismissed as experimental error. But, plotting *g* ratio as a relation of axon diameter (panel 1B), rather than fiber diameter (panel 1A), shows that variable slopes (i.e. negative versus positive) also can involve artifacts associated with the choice of *x*-axis variable. This unexpected instability raises concerns about the biological significance of the slope component in the regression fit. Concurrently, the *y*-intercepts are variable, which raises additional concerns. For example, should the average *g* ratio of the cohort be defined by the average of the *y*-intercepts because it is a common reference point (i.e. *x* = 0)? If so, the average *g* ratios from panels 1A and 1B would differ (0.84 versus 0.85; paired t-test, P = 0.007, *n* = 3). Alternatively, should the grand average defined in panel C be used? If so, how would this value be related to specific axon/fiber calibers when the regression slopes can change sign depending on the *x*-axis variable?

An additional important but subtle difference between panels 1A and 1B is the rank order of data points along the *x*-axis, which can change relative to nearby points. For example, within same-size rectangles, two of the three mouse A data points (green) in rectangle 1A appear to be left shifted in 1B, relative to the mouse B and C data points (red and black). Such shifts are data point-specific and nonuniform. A fourth mouse A data point left-shifts into the rectangle in panel 1B, and additional relative changes occur for all three mice. Thus, differences in rank order of the data points along the *x*-axis, and possibly other types of artifact, are present and may have substantial overall impact on the variability of the regression lines. Indeed, while compression of the range on the *x*-axis can increase regression slope (either more positive or more negative), it cannot change the sign as occurs for mouse C.

### A predictive axomyelin unit model emerging from first principles

Concerns raised about Fig. 1 necessitate a thorough examination of the raw experimental measurements, beginning with an exploratory data analysis (Filliben and Heckert, 2012) from the first principles – axon and fiber diameters. This critical first step is necessary to understand the relation between the two measured variables and identify artifacts contributing to regression fit instability. Thus, correlation plots of axon diameter as a relation of fiber diameter strongly suggests a linear relation, which is supported by extreme values of Pearson’s coefficients, *r_xy_*, indicating the data are nearly perfectly correlated in each mouse (Table 2). After pooling data from the three mice for convenience (Fig. 2A), the resulting line plot and *r_xy_* show the linear relation.

**Figure 2.**
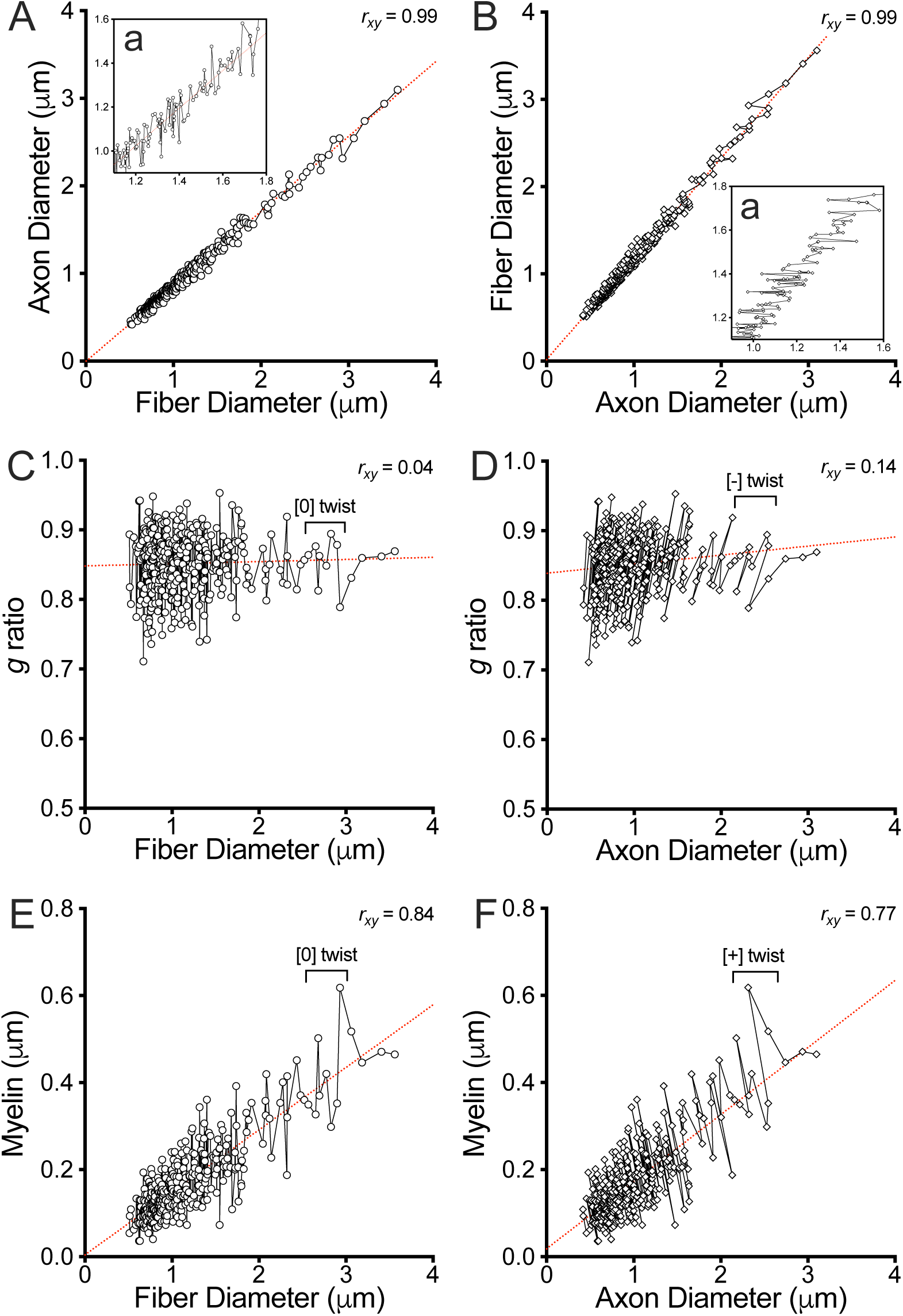
Correlation line plots reveal distortion artifacts in the data structure For ease of presentation, data from the three mice in Fig. 1 have been combined into a single dataset for exploratory data analysis. In all panels, data points are sorted by fiber diameter. **A.**, **B.** The near-perfect positive Pearson’s correlation coefficients strongly suggest a linear relation between axon and fiber diameters. Moreover, the regression lines pass through the Origin of the plots indicating the relation is directly proportional. Transposition of the axes in (A) (i.e. swapping the axes) changes the regression slope in (B) (Table 2) but not the Pearson’s coefficient. The predominantly vertical pattern of the point-to-point line in (A), insert a, is designated as zero twist ([0] twist). The point-to-point line is reflected across the regression line by the transposition in (B), inset a, yielding a data structure orthogonal to [0] twist. **C., D.** *g* ratio plotted as a relation of fiber or axon diameter, respectively. Pearson’s correlation coefficients are weak (Table 3). The data structure in (C) exhibits [0] twist, but is rotated clockwise in (D) ([-] twist). **E.**, **F.** plots show moderate Pearson’s correlations for myelin as a relation of fiber or axon diameter, respectively. In (E), the point-to-point line has [0] twist similar to panels (A) and (C). In contrast, the structure in (D) is rotated counterclockwise ([+] twist). Thus, plots in which fiber diameter is the independent variable maintain a consistent point-to-point line structure, while axon diameter as the independent variable causes unstable line structures and alters Pearson’s correlation coefficients.

**Table 2.**
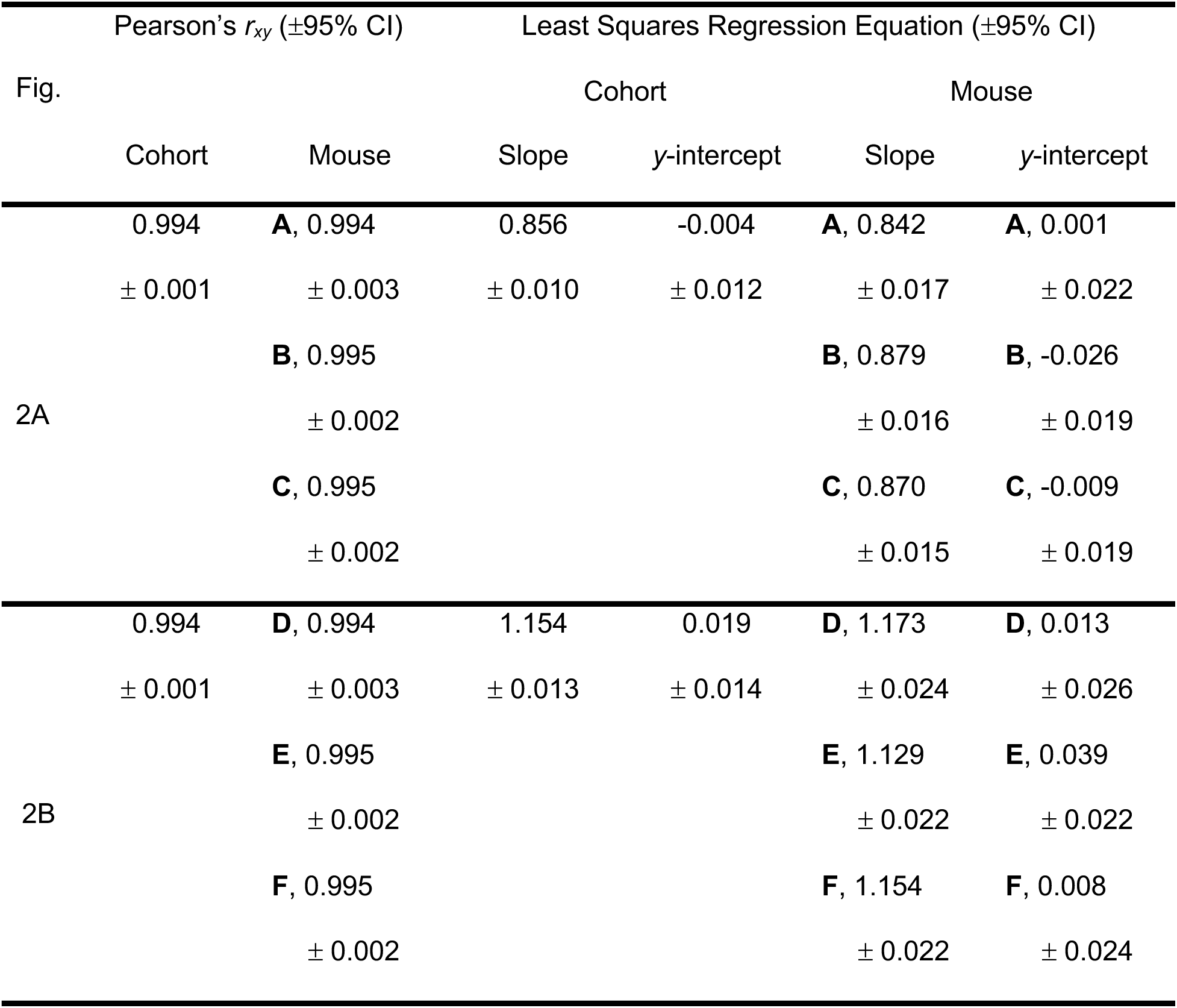
– axon diameter vs (fiber / axon diameter) plots for Fig. 2A and 2B.

| Fig. | Pearson's $r_{xy}$ ($\pm 95\%$ CI) | | Least Squares Regression Equation ($\pm 95\%$ CI) | | | |
| --- | --- | --- | --- | --- | --- | --- |
|  |  |  | Cohort |  | Mouse |  |
|  | Cohort | Mouse | Slope | y-intercept | Slope | y-intercept |
| 2A | 0.994 | **A**, 0.994 | 0.856 | -0.004 | **A**, 0.842 | **A**, 0.001 |
| | $\pm 0.001$ | $\pm 0.003$ | $\pm 0.010$ | $\pm 0.012$ | $\pm 0.017$ | $\pm 0.022$ |
|  |  | **B**, 0.995 |  |  | **B**, 0.879 | **B**, -0.026 |
| | | $\pm 0.002$ | | | $\pm 0.016$ | $\pm 0.019$ |
|  |  | **C**, 0.995 |  |  | **C**, 0.870 | **C**, -0.009 |
| | | $\pm 0.002$ | | | $\pm 0.015$ | $\pm 0.019$ |
| 2B | 0.994 | **D**, 0.994 | 1.154 | 0.019 | **D**, 1.173 | **D**, 0.013 |
| | $\pm 0.001$ | $\pm 0.003$ | $\pm 0.013$ | $\pm 0.014$ | $\pm 0.024$ | $\pm 0.026$ |
|  |  | **E**, 0.995 |  |  | **E**, 1.129 | **E**, 0.039 |
| | | $\pm 0.002$ | | | $\pm 0.022$ | $\pm 0.022$ |
|  |  | **F**, 0.995 |  |  | **F**, 1.154 | **F**, 0.008 |
| | | $\pm 0.002$ | | | $\pm 0.022$ | $\pm 0.024$ |

**Table 3.** – *g* ratio vs (fiber / axon diameter) plots for Fig. 2C and 2D.

| Fig. | Pearson's $r_{xy}$ ($\pm 95\%$ CI) | | Least Squares Regression Equation ($\pm 95\%$ CI) | | | |
| --- | --- | --- | --- | --- | --- | --- |
|  |  |  | Cohort |  | Mouse |  |
|  | Cohort | Mouse | Slope | *g* ratio * | Slope | *g* ratio * |
| 2C | 0.038 | **A**, 0.006 | 0.003 | 0.848 | **A**, 0.001 | **A**, 0.842 |
| | $\pm 0.104$ | $\pm 0.176$ | $\pm 0.008$ | $\pm 0.011$ | $\pm 0.015$ | $\pm 0.019$ |
|  |  | **B**, 0.140 |  |  | **B**, 0.011 | **B**, 0.840 |
| | | $\pm 0.183$ | | | $\pm 0.016$ | $\pm 0.019$ |
|  |  | **C**, -0.013 |  |  | **C**, -0.001 | **C**, 0.862 |
| | | $\pm 0.177$ | | | $\pm 0.014$ | $\pm 0.018$ |
| 2D | 0.138 | **D**, 0.111 | 0.013 | 0.839 | **D**, 0.011 | **D**, 0.832 |
| | $\pm 0.101$ | $\pm 0.170$ | $\pm 0.010$ | $\pm 0.011$ | $\pm 0.017$ | $\pm 0.019$ |
|  |  | **E**, 0.232 |  |  | **E**, 0.021 | **E**, 0.832 |
| | | $\pm 0.174$ | | | $\pm 0.017$ | $\pm 0.018$ |
|  |  | **F**, 0.082 |  |  | **F**, 0.007 | **F**, 0.854 |
| | | $\pm 0.178$ | | | $\pm 0.016$ | $\pm 0.018$ |

\*, the *g* ratio is the *y*-intercept of the least squares regression equation

In such light, linearity of the axon-fiber diameter relation is reasonably assumed to be a biological property of the myelin internode. This assumption establishes a statistical model, called the axomyelin unit model, which is valuable because of its predictive capacity. For example, the assumption justifies fitting a linear regression model to the axon-fiber diameter plot (red dotted line Fig. 2A, Table 2) so that a specified fiber diameter provides an estimate of the corresponding axon diameter. Extrapolation of the regression fit shows it passes through the Origin (Table 2, *y*-intercept = −0.004, 95% CI: [-0.017, 0.008]), demonstrating the axon-fiber diameter relation is directly proportional.

Such predictions accord with myelin internode biology. For example, the smallest myelinated axons have very thin myelin sheaths (1 or 2 wraps), so axon diameter ≅ fiber diameter and a linear regression fit should pass through, or very close to the Origin (within experimental error). Indeed, several statistical tests and biological constraints demonstrate this so-called direct proportionality property of the axon-fiber diameter relation (supplemental results, Fig. S1). In addition, the axomyelin unit model concurs with early results (Donaldson and Hoke, 1905) of the approximately linear relation between axon and myelin cross-sectional area in the PNS.

### Plots of data structure are distorted by the choice of independent variable

The data in Fig. 2A are sorted according to fiber diameter, and adjacent data points are connected by line segments (solid black). Zooming in (inset a), the plot reveals a persistent structure reasonably described as a predominantly vertical-pattern parallel to the *y*-axis. For purposes detailed below, the vertical pattern is designated as “zero twist” ([0] twist). Figure 2B shows the same data from Fig. 2A, but the axes are reversed (the data remain sorted by fiber diameter). Now the structure pattern is orthogonal to the *y*-axis (inset a), as expected.

In panels 2C and 2D, pooled *g* ratios are plotted as relations of fiber and axon diameter, respectively. Although the data in these panels are similar overall, there is an obvious clockwise rotation of the data in panel 2D with respect to the regression fit, which we designate as a negative twist ([-] twist) artifact. The origin of the artifact is unclear; however, the magnitude of the twist appears to increase in proportion to the axon diameter. Accompanying the twist, the data points on the right of the plot are increasingly asymmetric across the dotted line, suggesting the regression fit may be of lower quality than in panel 2C.

The Pearson’s coefficient in panel 2C is very weak and the 95% CI includes zero (i.e. a horizontal regression fit), as does the regression slope (Table 3). Indeed, the standard equation for a linear regression fit (*y* = slope * *x* + *y*-intercept) is reduced to a constant value (the *y*-intercept) when the slope = 0, so *g* ratio is invariant (uncorrelated) with fiber diameter, by definition. This result is also a prediction from the direct proportionality assumption of the axomyelin unit model.

Pearson’s coefficient from panel 2D is much larger than panel 2C, which suggests *g* ratio and axon diameter are correlated. Consistent with this notion, both the Pearson’s coefficient and the slope are statistically non-zero (Table 3, the 95% CIs do not bracket zero). Indeed, paired t-tests of individual *r_xy_* values and slopes between panels 2C and 2D are statistically significant (P = 0.001 and P = 0.005, respectively), lending support to dependence of the *g* ratio on axon diameter. However, such dependence is precluded because of the direct proportionality of the axomyelin unit model, where *g* ratio is predicted to be constant. Thus, the choice of axon diameter as the independent variable may exacerbate experimental error in the raw data leading to an artifact. Presumably the larger error is associated with range compression of the *x*-axis and/or local reordering of the data points, which may generate the [-] twist artifact.

Panels 2E and 2F show correlations for myelin diameter as a relation of fiber or axon diameter, respectively. The Pearson’s coefficients for the pooled and individual data are of moderate strength (Table 4) and far weaker than in panels 2A and 2B, suggesting increased experimental error as a result of computing myelin diameter (2 x myelin radial thickness) by subtracting axon diameters from fiber diameters. The Pearson’s coefficient from panel 2F is also markedly lower than panel 2E, despite the simple substitution of axon for fiber diameter as the independent variable. This difference is statistically significant (paired t-test, P = 0.004). Importantly, the regression fit in panel 2E passes through the Origin (Table 4), as predicted by the axomyelin unit model, but this is not the case for panel 2F where the regression fit implies the impossible, that axon diameter exceeds fiber diameter.

**Table 4.** – myelin vs (fiber / axon diameter) plots for Fig. 2E and 2F.

| Fig. | Pearson's $r_{xy}$ ($\pm 95\%$ CI) | | Least Squares Regression Equation ($\pm 95\%$ CI) | | | |
| --- | --- | --- | --- | --- | --- | --- |
|  |  |  | Cohort |  | Mouse |  |
|  | Cohort | Mouse | Slope | y-intercept | Slope | y-intercept |
| 2E | 0.839 | **A**, 0.857 | 0.144 | 0.004 | **A**, 0.158 | **A**, -0.001 |
| | $\pm 0.036$ | $\pm 0.041$ | $\pm 0.010$ | $\pm 0.012$ | $\pm 0.017$ | $\pm 0.022$ |
|  |  | **B**, 0.815 |  |  | **B**, 0.123 | **B**, 0.025 |
| | | $\pm 0.055$ | | | $\pm 0.017$ | $\pm 0.020$ |
|  |  | **C**, 0.849 |  |  | **C**, 0.142 | **C**, -0.004 |
| | | $\pm 0.043$ | | | $\pm 0.016$ | $\pm 0.021$ |
| 2F | 0.775 | **D**, 0.794 | 0.154 | 0.019 | **D**, 0.173 | **D**, 0.013 |
| | $\pm 0.039$ | $\pm 0.057$ | $\pm 0.013$ | $\pm 0.014$ | $\pm 0.024$ | $\pm 0.026$ |
|  |  | **E**, 0.753 |  |  | **E**, 0.129 | **E**, 0.039 |
| | | $\pm 0.072$ | | | $\pm 0.022$ | $\pm 0.022$ |
|  |  | **F**, 0.790 |  |  | **F**, 0.154 | **F**, 0.008 |
| | | $\pm 0.059$ | | | $\pm 0.022$ | $\pm 0.024$ |

A reasonable explanation for moderate correlations in panels 2E and 2F is compounding of experimental errors by the subtraction operation to compute myelin diameter (experimental errors are additive when measured variables are added or subtracted). Nevertheless, choosing axon diameter as the independent variable appears to exacerbate the impact of these errors. Perhaps the [+] twist in panel 2F relative to panel 2E, further reduces the correlation. Although the origins of these artifacts are unclear, they indicate that using axon diameter as the independent variable yields data structure distortions that affect the properties of the data structure in multiple ways.

### Simulations reveal the origin of artifacts in *g* ratio plots

Differences in Pearson’s coefficients, linear regression slopes and *y*-intercepts in Figs 1 and 2 are, at least in part, associated with axon diameter as the independent variable. In contrast, fiber diameter plots accord with axon-fiber diameter direct proportionality, but other unknown errors that cannot not be discounted. To identify such unknowns, simulations can be used to explore artifacts in the absence of experimental errors.

In Fig. 3A, the green simulated values (*g* ratio = 0.64, designated as a *g* ratio set) are plotted as a relation of fiber diameter and constitute a horizontal row. The largest fiber, 18.8μm, has axon diameter = 12μm. Three other *g* ratio sets (*g* ratios = 0.7, dark blue; 0.76, light blue; and 0.82, red) are also plotted. The largest fiber of the red *g* ratio, also 18.8μm, has a significantly larger axon diameter (15.4μm) because the myelin sheath is thinner (*g* ratio = 0.82 versus 0.64). A regression fit to these values has zero slope (= 10^-17^), and the *y*-intercept equals the grand average *g* ratio because of the evenly spaced *g* ratio set distributions on both the *x*- and *y*-axes. Figure 3B shows the same simulated *g* ratios as panel 3A, but plotted as a relation of axon diameter. Again, the *g* ratio sets form horizontal rows, but range compression of the *x*-axis (analogous to Fig. 1B) is *g* ratio-dependent, so the green set is compressed to a greater extent than the red set. This distortion accounts for the different order of data points along the *x*-axis in Fig. 1A versus 1B (rectangles). In addition, the regression fit has a positive slope and the *y*-intercept is lower than the grand average *g* ratio in panel 3A.

**Figure 3.**
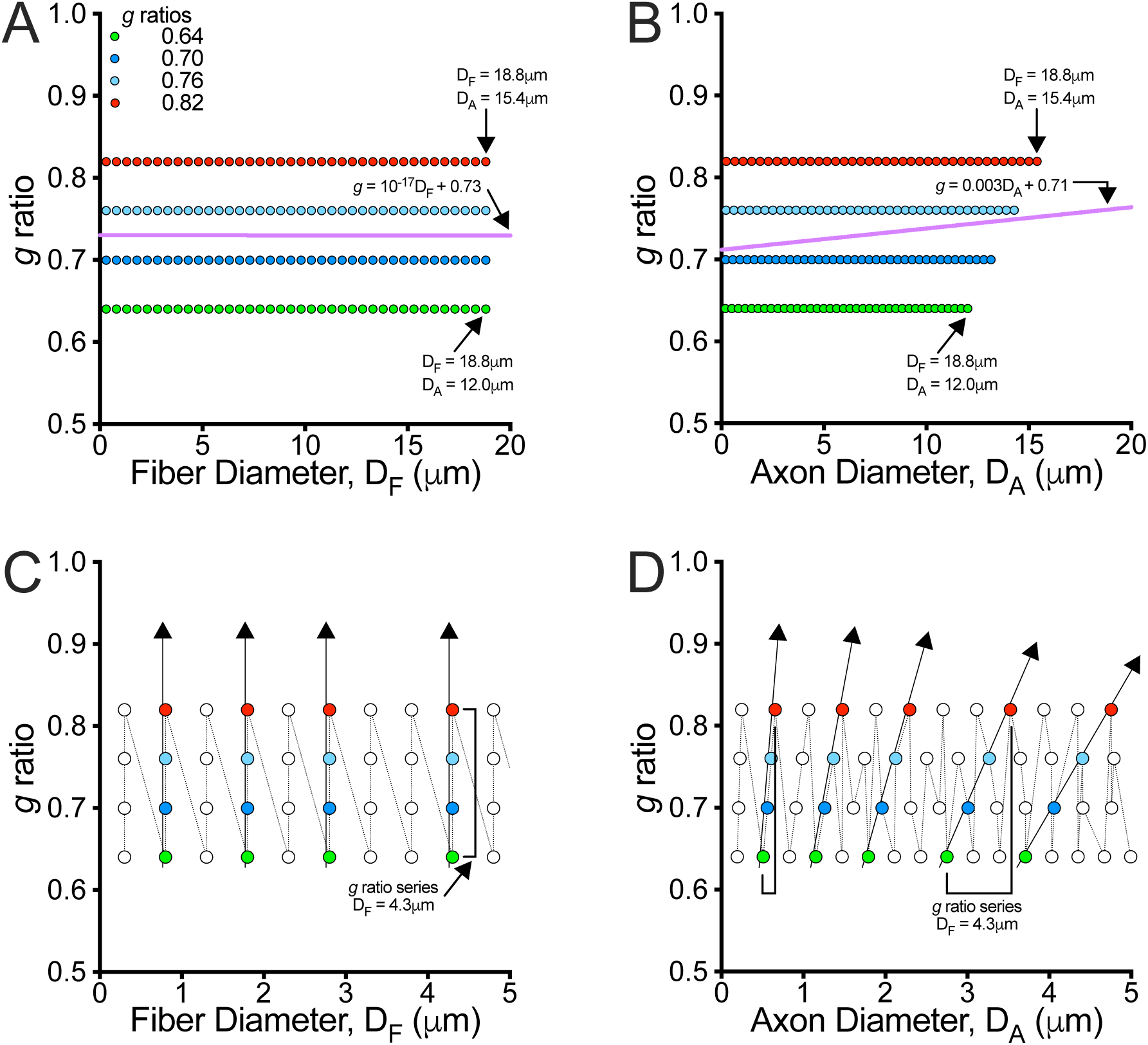
Distortion artifacts in *g* ratio simulations Simulations of evenly spaced *g* ratio values {.64, .70, .76 and .82} plotted as relations of fiber diameter, with like-colored values designated as *g* ratio sets (horizontal rows). **A.** *g* ratio plotted as a relation of fiber diameter (.3-18.8μm). The regression fit has zero slope and the grand average *g* ratio = .73. **B.** *g* ratio as a relation of axon diameter (.18-15.4μm.). Compression of the *x*-axis range (analogous to Fig. 1), is actually *g* ratio-dependent, where the green *g* ratio set is compressed to a greater extent than the red set. This distortion generates an uneven data structure (right hand side), a positive slope in the regression fit and decreases the *y*-intercept. **C.** Expanded view of (A) shows *g* ratios organized in ten series of four points. Four are colored to emphasize their ascending vertical arrangement (arrows, .64 → .82). The balanced structure and order (connecting dotted line) extends along both the *x*- and *y*-axes. **D.** Sequential *g* ratios in each series are not organized vertically and the connecting line does not necessarily join consecutive points within a series.

Zooming in on panel 3A (panel 3C) shows the *g* ratio values are organized into multiple *g* ratio series comprising four points parallel to the *y*-axis. Four of these series are colored by *g* ratio to emphasize their ascending vertical organization (arrows, 0.64 → 0.82). Such a balanced structure and order extends along both the *x*- and *y*-axes and is reminiscent of the data structure in Fig. 2C. In contrast, *g* ratio structure breaks down when plotted against axon diameter in panel 3D. Small axons have a consistent *g* ratio order but sequential *g* ratios in each series have increasing axon diameters so the arrows exhibit [-] twist which intensifies as axon diameter increases. Further, for larger diameter axons the point-to-point line does not even join consecutive points within a *g* ratio series. This distorted and disorganized structure may account for both the positive regression slopes and [-] twist artifact in Fig. 2D and panels 3B and 3D.

### Random sampling contributes to regression slope instability in *g* ratio plots

Simulations in Fig. 3 reveal a regression slope artifact when axon diameter is used as the independent variable. In contrast, this artifact is absent in fiber diameter plots and the regression fit has zero slope. These differences notwithstanding, a slope artifact is associated with random sampling, which is typical methodology in real experiments. For example, a typical experiment in optic nerve involves measuring a small number of *g* ratios (typically 100-200) from 60,000 ± 8,000 fibers depending on mouse strain (Williams et al., 1996; Seecharan et al., 2003). Figure 4 shows three random samples taken from simulated *g* ratios in Fig. 3. For each random sample, the RAND() function in Excel (Microsoft, ver 16.66.1) was used to select 30% of the simulated *g* ratios {0.64, 0.70, 0.76 and 0.82} for plotting as relations of fiber diameter (panels 4A, C and E) or axon diameter (panels 4B, D and F). Together, the panels demonstrate that regression slopes of random samples can be positive (panels 4A, B), flat (panels 4C, D) or negative (panels 4E, F), depending on the random combination of *g* ratios and the choice of independent variable. Presumably, this type of artifact may contribute to experimental errors in all *g* ratio plots, and in similar fashion to Fig. 1, *x*-axis compression alters regression slopes and *y*-intercepts. This, again, raises concern about the biological relevance of simple linear regression fits for the analysis and interpretation of *g* ratio plots.

**Figure 4.**
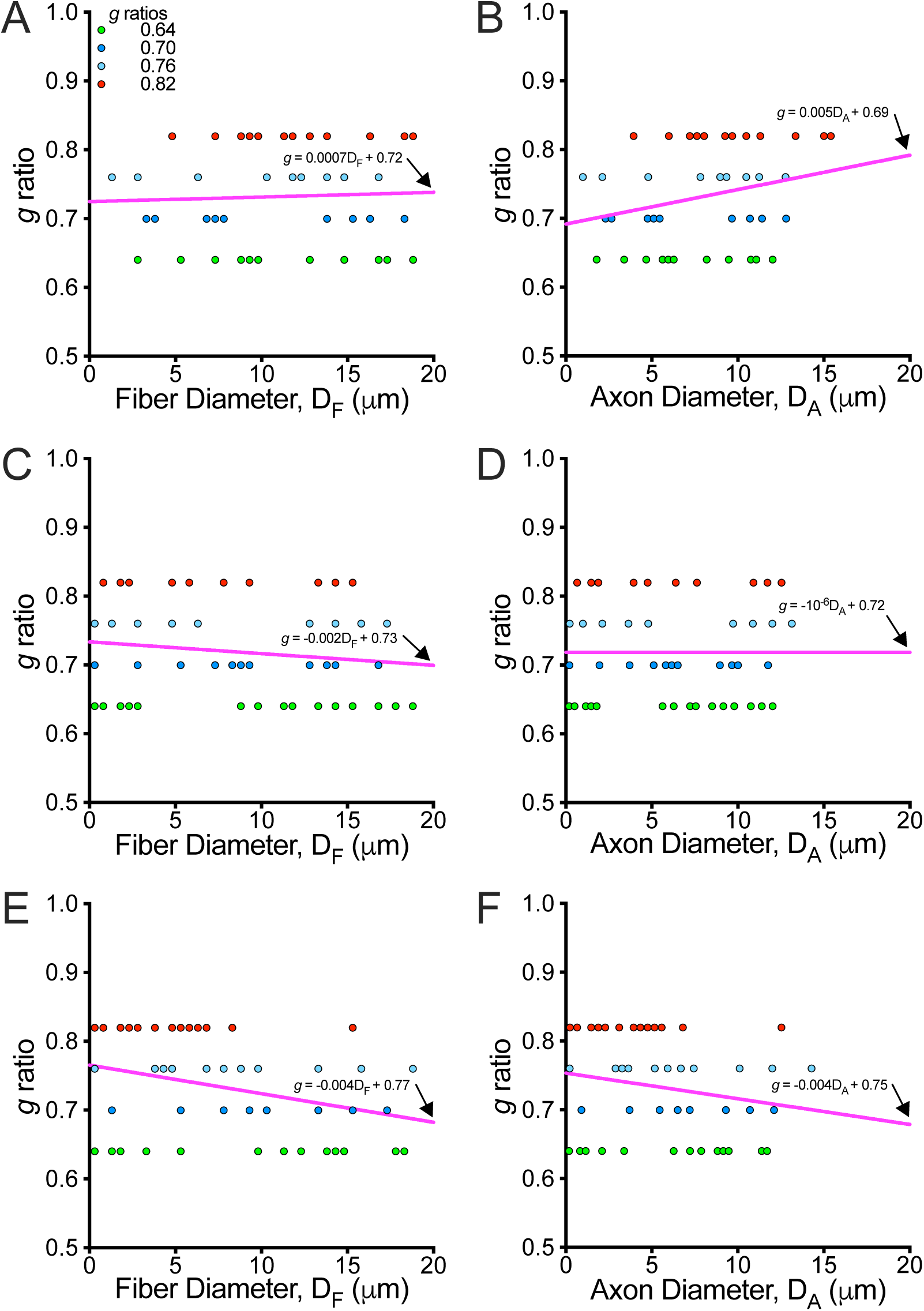
Random sampling artifacts cause variable regression slopes in all *g* ratio plots **A., C., E.** Simulated *g* ratio sets from Fig. 3A were randomly sampled (∼30% of the values), plotted as relations of fiber diameter and fit by linear regression. The regression slopes are stochastic (positive or negative) because they depend on the combination of values sampled, rather than maintaining the zero slope of the parent set. **B., D., E.** The randomly sampled simulated values from (A, C, E), replotted as relations of axon diameter, also have stochastic slopes. This random sampling artifact affects all *g* ratio plots but can be additive with other artifacts like *x*-axis compression (Fig. 3B).

### Simulations reveal artifacts in myelin plots

In addition to artifacts in *g* ratio versus axon diameter plots (Figs 2D, 3B and D), the same is observed in myelin versus axon diameter plots (Fig. 2F). The simulated values generated for Fig. 3 have been used to derive myelin diameter plots (myelin diameter = fiber diameter – axon diameter) in Fig. 5. Panel 5A shows an even distribution of values along both axes, although spacing within the red set is marginally closer compared to the green set (and the blue sets). The spacing differences notwithstanding, the regression fit passes through the Origin symmetrically bisecting the plot, and the largest fibers are vertically organized in decreasing myelin diameter (equivalent to *g* ratios 0.64 → 0.82 in Fig. 3) to yield a balanced data structure.

**Figure 5.**
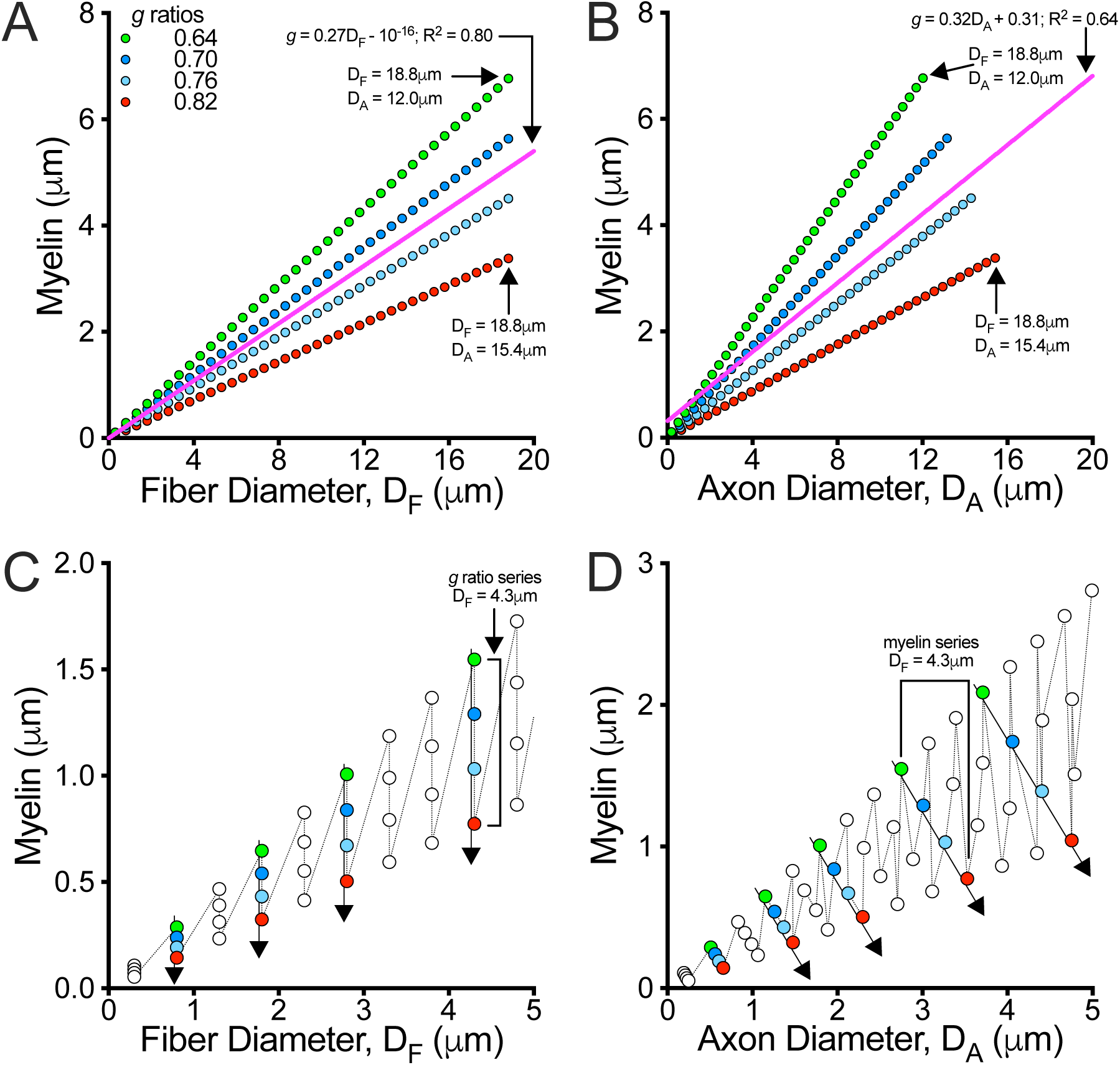
Distortion artifacts in simulated myelin diameter plots **A.** Simulations of myelin diameters derived from the *g* ratio values in Fig. 3, plotted as a relation of fiber diameter. The like-colored sets form diagonals with *g* ratio-dependent slopes that are inversely-related to myelin diameter and the largest fibers are vertically aligned (similar to Fig. 3A). The regression curve falls along the line of symmetry of the plot and passes through the Origin, indicative of direct proportionality. **B.** The same myelin diameters as in (A) but plotted as a relation of axon diameter. The *x*-axis compression is myelin diameter-dependent, with the green set compressed to a greater extent than the red (similar to Fig. 3B). The regression curve lies off the line of symmetry for the plot and does not pass through the Origin. **C.** Expanded view of (A) shows myelin diameters organized in ten series of four points. Four are colored to emphasize their descending vertical arrangement (arrows, .64 → .82). The balanced structure and order (connecting dotted line) is apparent along both the *x*- and *y*-axes. **D.** Sequential myelin diameters in each series are not organized vertically, and the connecting line does not necessarily join consecutive points within a series.

The myelin versus axon diameter plot in panel 5B is overall similar to panel 5A (e.g. spacing within the red set is marginally closer than the other sets) but with several significant differences. First, the regression fit divides the plot asymmetrically, suggesting the structure of the simulated values is distorted. In addition, the fit poorly represents small diameter axons and does not pass through the Origin. Indeed, the *y*-intercept indicates axons with zero diameter are myelinated. Second, there is a myelin diameter-dependent compression along the *x*-axis so that axons are more closely spaced, and the largest axon is larger, for the red set compared to the other sets. This distortion is distinct from that in Fig. 3B, possibly because myelin diameter and *g* ratio are inversely-related (i.e. thicker myelin = lower *g* ratio).

In Fig. 5C, which zooms in on panel 5A, the familiar vertical and organized structure of fiber diameter plots is apparent and analogous to Fig. 3C except the *g* ratio series arrows are downward-directed. In contrast, the twist structure in panel 5D is again oblique to the axes, but in the opposite direction to Fig. 3D, downward-directed with an axon diameter-independent twist. This distortion is similar to Fig. 2F. The inverse proportionality of myelin diameter and *g* ratio notwithstanding, the clockwise (Fig. 2D) and counterclockwise twists (Fig. 2F) also may not be entirely analogous because of their distinct twist properties (for *g* ratio, twist is axon diameter-dependent; for myelin diameter, twist is axon diameter-independent).

### Artifacts contribute to nonlinear relations between myelin and axon diameter

In 1947, Sanders (1947) published a large influential study purporting to demonstrate that neither *g* ratio nor myelin diameter maintain constant proportionality as axon diameter increases. We now know that a number of the conclusions were misinterpreted because of technical artifacts. For example, reliance on small caliber axon measurements (< 3-4μm) that were below the resolving power of his light microscope (Schnepp and Schnepp, 1971; Price and Sprich, 1975), while for large caliber axons the regression fits were more consistent with a linear myelin versus axon diameter relation and constant *g* ratio (Sanders, 1947).

Nonetheless, as a follow up on the Sanders assertions, a random sample of the simulated values from Fig. 5 is plotted in Fig. 6A and 6B as a relation of fiber or axon diameter, respectively. Both plots are fit with linear and logarithmic regression models, to determine if curvilinear fits can be generated in the absence of experimental error, and corrected Akaike’s information criteria (AICc) tests (Dziak et al., 2020) are used to assess which fit is more likely to be correct. In panel 6A, both model fits to the myelin versus fiber diameter relation appear to be linear. And although the linear regression fit includes the Origin (*y*-intercept = 0.189, 95% CI: [-0.171, 0.550]), the confidence intervals for the logarithmic fit are very wide and unstable. The AICc test indicates there is little evidence to favor either model, with the probability that the linear fit is correct being AICc = 52% compared to AICc = 48% for the logarithmic fit.

**Figure 6.**
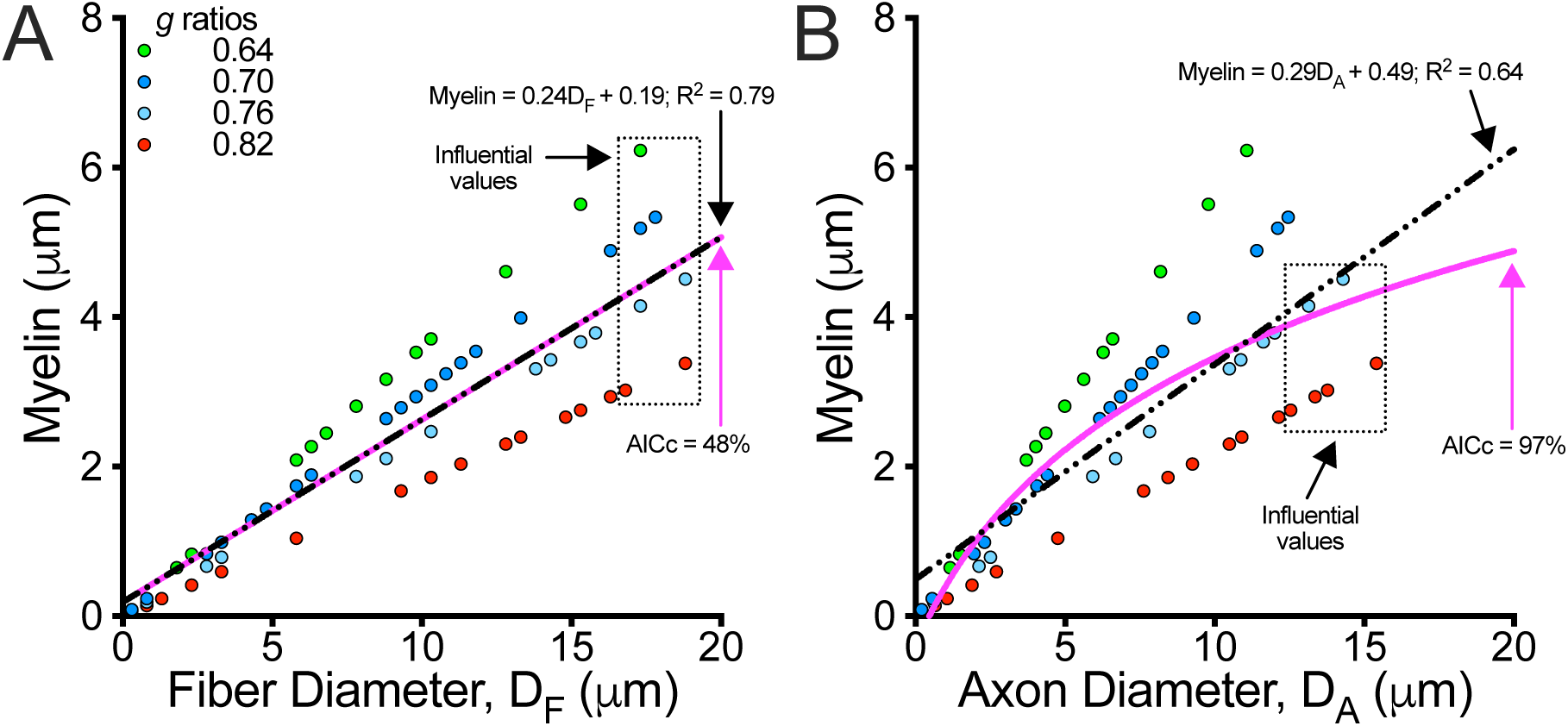
Simulated myelin diameters plotted against fiber and axon diameters are linear Randomly sampled myelin diameters from Fig. 5A are moderately linearly correlated to fiber or axon diameter. **A.** Values plotted as a relation of fiber diameter and fit with linear and logarithmic regression models. The model fits are compared using Akaike’s Information Criterion (AICc) and show equal probabilities of correctness for a linear fit = 52% versus a logarithmic fit = 48%. **B.** Values from (A) plotted as a relation of axon diameter and fit with linear and logarithmic regressions. In contrast to (A), AICc = 3% for a linear fit compared to AICc = 97% for a logarithmic fit. However, the nonlinear fit is likely an artifact associated with the 6 largest values (dotted rectangle), which are asymmetric across the regression line. Indeed, without these strongly influential values, AICc = 48% for a linear fit, while removing any other six sequential points marginally changes the logarithmic fit preference (AICc = 1-11% (median = 5%) for a linear fit). In (A), there is weak evidence that removing the fibers in the rectangle alters the preference for the fit, where AICc = 12-19% (median = 18%) for a linear fit. But removing any other six fibers has minimal impact, where AICc = 44-68% (median = 57%) for a linear fit.

In contrast, panel 6B shows the model fit to the myelin versus axon diameter relation is likely logarithmic, with AICc = 97%. However, detailed examination of the data structure shows that most of the largest axons in panel 6B (dotted rectangle) are asymmetric across the linear regression curve (the same fibers in panel 6A are symmetric). This diminishes the fit and drives the nonlinear shape of the curve. Indeed, removing the asymmetric points renders the linear and logarithmic fits indistinguishable, with AICc = 48% probability for a linear fit that passes through the Origin. On the other hand, removing any other six consecutive points in the graph does not change the high AICc probability of a logarithmic fit. Thus, the points in the rectangle have an unduly strong effect on the model (i.e. influential points).

Asymmetric distortions in data structure can also occur near the Origin, for example because of systematic limit of resolution measurement errors. These errors were very common prior to use of the electron microscope in myelin research (Schnepp and Schnepp, 1971; Price and Sprich, 1975), and even persist in the current literature when the pixel resolution of micrographs is too low (i.e. >3nm). Accordingly, Sanders’s (1947) assertion that a nonlinear myelin-axon diameter relation exists in his dataset is readily explained by light microscopy limit of resolution errors (known as the Rayleigh criterion). Indeed, he acknowledged at the time that his measurements of the smallest axons were asymmetrically distributed.

### Measurement error artifacts in *g* ratio plots

There is lingering uncertainty associated with the variable Pearson’s coefficients in Fig. 2 because the same axon-fiber diameter data pairs are used in all six panels with only the simple arithmetic manipulations, subtraction and division. But this variation can be explained by the impact of those manipulations, which compound random experimental errors. Thus, for addition/subtraction (panels 2E and F), absolute errors of each variable are additive; for multiplication/division (panels 2C and D), the percentage errors are additive.

In Fig. 7A a simulated errorless set of axon diameters between 0.18-4μm with myelin diameter corresponding to a constant *g* ratio = 0.70 (from the dark blue set in Figs 3-6) is perfectly correlated as indicated by the Pearson’s coefficient and the regression fit through the Origin. The axon diameter residuals of the fit are plotted against fiber diameter and are all zero (red line, Fig. 7B). In panel 7C, the same axon and fiber diameters are transformed to introduce random errors of measurement between ±0.1μm (several times smaller than the optical resolution of the light microscope) and plotted with the regression fit (panel 7D, residuals are randomly and symmetrically distributed). The scatter in the plotted values is even and reminiscent of real data in Fig. 2A and the regression fit signifies direct proportionality of the axon-fiber diameter relation.

Panel 7E reveals the extent to which experimental errors are magnified by the *g* ratio transformation (eqn 1). Despite the even and randomly distributed residuals in panel 7D, the panel 7F residuals are nonuniform and widely dispersed near the *y*-axis, while being more constrained to the right. Thus, demonstrably uniform measurement errors (panel 7D) are dramatically distorted by the *g* ratio transformation. Moreover, the distortion drives the non-zero slope of the regression fit and is not compatible with direct proportionality shown in panel 7C. These distortion artifacts have such a strong effect because they violate the assumptions of linear regression, causing unstable positive or negative regression slopes, as observed in Fig. 1.

**Figure 7.**
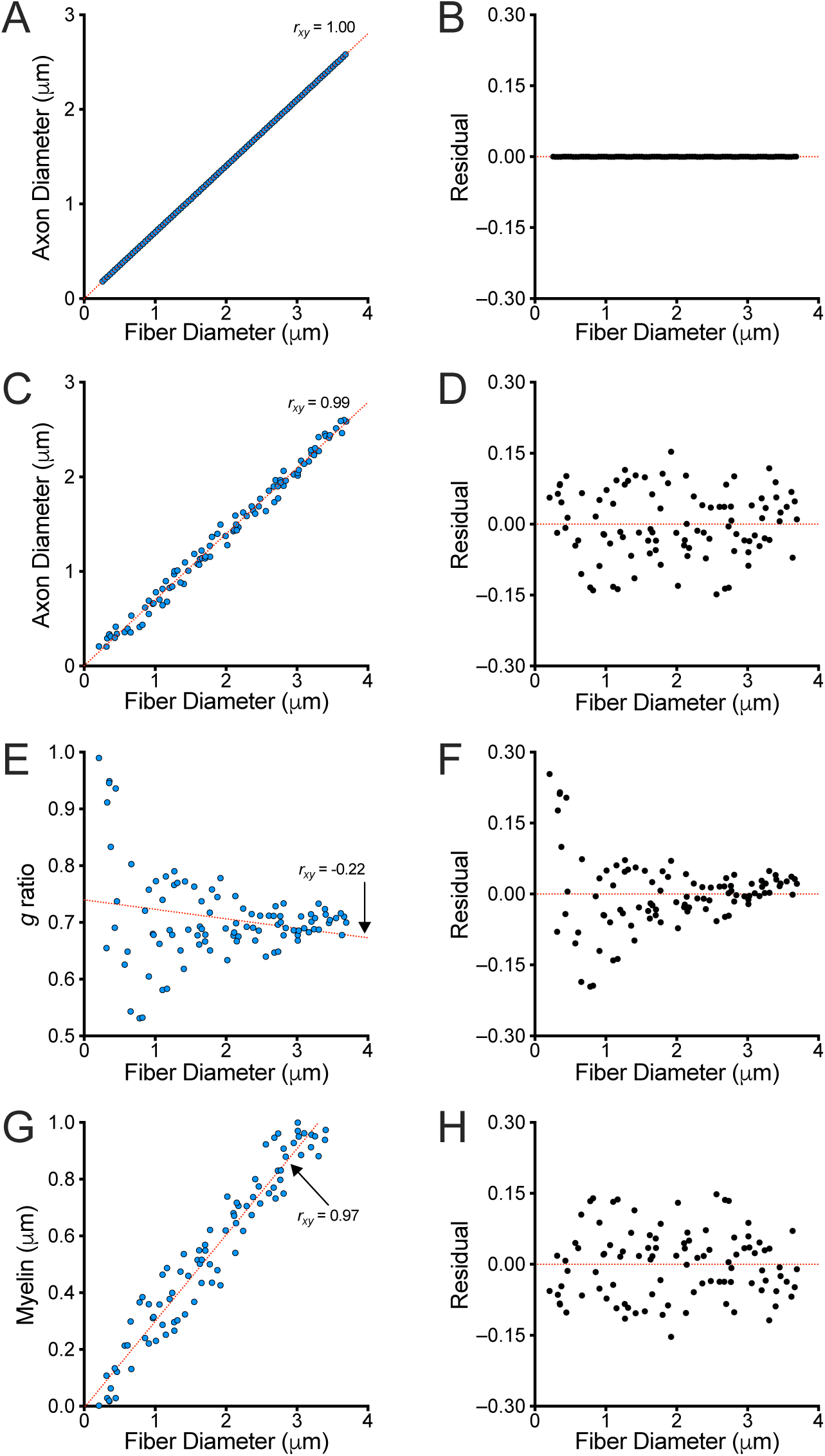
Simulations of axomyelin unit variables show linear regression of *g* ratio plots are unbalanced **A.** Errorless simulation plot of the axon-fiber diameter relation using a single value of *g* ratio = .70 to compute fiber diameters from a series of axon diameters that are incremented by .025μm over the range, .18 – 2.6μm. The Pearson’s coefficient, *r_xy_* = 1, is perfect, and a linear regression fit of the simulation passes through the Origin (red dotted line). **B.** A plot showing the distance (i.e. the linear regression residuals) of each axon diameter value in (A) from the linear regression line as a relation of fiber diameter. All the residuals = 0 because the absence of error means that all values in (A) are on the regression line. **C.** Introduction of random errors (Gaussian, S.D. = .1μm on the *x*- and *y*-axes) into the axon and fiber diameter values reduces *r_xy_* = .99. Importantly, the simulated values are randomly and evenly distributed across the regression line (red dotted line), so it passes through the Origin. **D.** The residuals plot from (C) shows the introduced errors are random (no patterns) and form a uniform band on either side of the red dotted line. **E.** The axon and fiber diameter values from (C) used to generate *g* ratios and plotted versus fiber diameter. The regression fit (red dotted line) has a negative slope, indicating that *g* ratio and fiber diameter are somewhat correlated (*r_xy_* = −.22). But this property is discordant with direct proportionality shown in (C) and must arise from artifact. **F.** Residuals from the plot in (E) no longer form a uniform band across the red dotted line, indicative of a poor regression fit or uncontrolled errors. The regression fit poorly summarizes the plot. **G.** Myelin versus fiber diameter plot from the simulation values in (C). The values are random and uniformly distributed across the regression fit. The Pearson’s coefficient *r_xy_* is marginally lower than the corresponding plot in (C). **H.** The residual plot for the values in (G) shows the errors are similar to the plot in (D).

Finally, the organization of values in the myelin verses fiber diameter plot (panel 7G) shows that the errors associated with the subtraction transformation are modest and the organization of the values is similar to panel 7C, and the Pearson’s coefficient is only marginally lower. As asides, the similar organization of simulated values in panels 7E and 7F is not coincidental; the regression slope in panel 7E is close to the horizontal slope of the red dotted line in panel 7F. Also, the residual distributions in panels 7D and 7H are uniform, and perfect reflections across the red horizontal line).

## DISCUSSION

The exploratory data analysis used in this study comports with the widely known property of myelinated axons, that of an extremely strong linear correlation between axon and fiber diameters measured in electron micrographs of white matter tracts from many species. But detailed statistical analysis demonstrates that the axon-fiber diameter relation is directly proportional (Fig. S1), an apparently novel perspective that has significant implications for *g* ratio analysis and interpretation. Moreover, formal adoption of this direct proportionality relation establishes an explicit model of the axomyelin unit under physiological conditions. The importance of such an approach cannot be overstated because it can be used to predict axomyelin properties and minimize ad hoc or inappropriate analyses and interpretations in favor of constrained and rigorous hypothesis development and testing.

Defining a linear axomyelin unit model, using experimental data under physiological conditions, and simulations, is a powerful approach. The model accords with previous seminal studies in the peripheral and central nervous systems (Donaldson and Hoke, 1905; Rushton, 1951; Schnepp and Schnepp, 1971; Waxman and Bennett, 1972; Price and Sprich, 1975). In addition, it is consistent with expected properties of myelin internodes: (1) diameters of unmyelinated axons are equal to their fiber diameters, so linear regression fits of axon versus fiber diameter plots pass through or very close to the Origin (Figs 2A, 2B, S1, Table 2) i.e. *x*-intercept ≥ 0, *y*-intercept ≤ 0, (2) the smallest axons have very thin myelin, so regression fits of myelin versus fiber diameter plots pass through or very close to the Origin (Fig. 2E, Table 4), (3) *g* ratios are not correlated with fiber diameter, so regression fits of *g* ratio versus fiber diameter plots have slope = 0 (Fig. 2C, Table 3). Importantly, these properties are more consistently demonstrated in graphs when fiber diameter is chosen as the independent variable.

A potential caveat of all myelin experiments (Fig. 2A) is the measurements themselves, because they constitute pseudo-replicate data (i.e. the measurement of multiple myelin internodes per mouse). While correlations and regression fits of data within a single mouse are often valid, most statistical tests comparing mice in a cohort or between treatment groups rely on every data point being uncorrelated to all other data points (i.e. a single independent measurement per mouse). Violations of this independence assumption can invalidate the conclusions from inferential statistical tests like t-tests and ordinary ANOVA. Mixed-effects analysis accounts for the systematic error effects of pseudo-replicate data and is increasingly available, but is also more difficult to interpret. The current study and the following article strike a balance to maximize accuracy and minimize interpretation difficulty.

### Artifacts in *g* ratio plots generated without prior exploratory data analysis

Detailed exploration of correlation plots from the Sham cohort (Figs 1 and 2) raises several potentially troubling issues for data analysis and assessment of myelin internodal properties in the contemporary literature. Not surprisingly, these issues will have significant implications for interpreting and understanding normal physiology, disease and dynamical states, and the success of remediation strategies for potential bioactive molecules. Further, conclusions drawn from *g* ratio plots in other emerging areas, including myelin plasticity and computer-automated *g* ratio measurements, may be tenuous in at least some instances. In this light, there is urgent need to standardize the design and implementation methodology for *g* ratio analyses. This topic is addressed in the accompanying article. In the current study, several problems associated with artifacts in *g* ratio plots have been identified.

The origins of some artifacts can be traced to distortions in data structure, which are revealed in line plots. For example, when fiber diameters are used to sort the data in ascending order, and plots are generated using axon diameter as the independent variable (Fig. 2D and 2F), the distortions are apparent as twist artifacts. Conversely, if axon diameters are used to sort the data, plots with fiber diameter as the independent variable have twist artifacts (not shown). Thus, the differences between plots against fiber or axon diameter are real so the important question becomes, which of these independent variables is appropriate/correct for summarizing myelin internode data?

In answer to this question, we can be certain that axon diameter as the independent variable causes/exacerbates the artifacts. The evidence is the statistically significant correlation between *g* ratio and axon diameter in Fig. 2D, which is incompatible with the direct proportionality of the axomyelin unit model. Further, two crucial assumptions of linear regression are not met in axon diameter plots – data should be uniformly distributed along the *x*-axis and residuals should be normally distributed about the regression fit. These distortions are minimized when fiber diameter is the independent variable.

On the other hand, an unexpected regression slope artifact is revealed in Fig. 4, which impacts *g* ratios plotted against either fiber or axon diameter. Similar to Figs 3 and 5, this artifact is associated with violating the assumptions of linear regression (non-uniform density of values along the *x*-axis) and stems from a universal experimental methodology in *g* ratio studies, that of randomly sampling a small number of myelinated fibers from a large nonuniform population. Several recent studies have sought to eliminate this issue using automated sampling of large numbers of myelinated fibers (Stikov et al., 2015; Janjic et al., 2019; Oost et al., 2023). However, a different artifact emerges, that of insufficient measurement resolution (the Rayleigh criterion) so that myelin around small axons cannot be accurately resolved and myelin sheath thickness is overestimated.

Early studies were also marred by this type of technical artifact, in particular the limit of resolution of light microscopy (either histology or birefringence methods). The deficiency is readily apparent from erroneous estimates of the smallest myelinated axons in the PNS – 2μm diameter – which is twice the actual lower threshold of myelinated axons. The limit of resolution issue was formally brought to light by theoretical considerations and systematic experimental methodologies in subsequent studies (Rushton, 1951; Schnepp and Schnepp, 1971; Price and Sprich, 1975; Waxman and Swadlow, 1976).

Although not the first to weigh-in on issues confronting the interpretation of *g* ratio plots, the pioneering and influential work of several groups (Duncan, 1934; Schmitt and Bear, 1937; Gasser and Grundfest, 1939; Taylor, 1942; Sanders, 1947) has significantly shaped the debate for almost a century and promulgated the view that *g* ratios plotted against axon diameters reveals important connections between internodal structure and function. However, Fig. 6 brings to light data structure artifacts – in the form of influential values at the left and right extremes of the *x*-axis range – that may account for nonlinear plots from experimental observations.

In light of Figs 2-7, we may be left with the perception that *g* ratio plots as relations of fiber or axon diameter are inexorably confounded by artifacts. These artifacts are likely additive, scale with the extent of data structure distortion and may be exacerbated as the range of *g* ratios increases (e.g. in non-physiological states). Nevertheless, there appear to be fewer sources of artifact (e.g. random sampling errors) in *g* ratio plots when a zero regression slope is used and fiber diameter is the independent variable.

With due consideration of the aforementioned concerns and statistical results, ultimately we must come back to the overarching question – what about the biological relevance? For example, the regression equations in Fig. 1 show the slopes are close to zero, even if statistically significant. Some artifacts in Figs 2-5 are also quite small, even if they are additive. Considering the steepest regression slope in Fig. 1A for mouse C (slope = 0.031), the smallest axon diameter = 0.37μm has *g* ratio = 0.74, while the largest axon diameter = 2.9μm has *g* ratio = 0.82. These *g* ratios deviate from the grand average (= 0.76, Fig. 1C) by 0.02 and 0.06, respectively, reflecting < 8% differences, which might reasonably be considered within the margin of error. Thus, notions of a fiber diameter-dependence in *g* ratios (or myelin diameter) in this cohort is moot, even though a regression slope is measurable.

In summary, the use of exploratory data analysis demonstrates that the relation between axon and fiber diameters is most likely directly proportional. There is no statistical evidence to the contrary. Thus, a predictive axomyelin unit model naturally emerges. This model imposes important constraints for correctly interpreting *g* ratio plots, which should be generated using fiber diameter as the independent variable to minimize artifacts and preserve Rushton’s (1951) theoretical predictions that observed differences in *g* ratios have minimal impact on the functional properties for fibers of the same caliber.

## SUPPLEMENTAL RESULTS

From an overall perspective in characterizing the axomyelin unit under physiological conditions, the importance of the axon-fiber diameter relation is primary. With a detailed understanding of this relation, it is possible to establish a predictive model of internodal myelin biology that can be used to formulate and test hypotheses. To this end, acceptance of the central assumption of direct proportionality based solely on Pearson’s and/or Spearman’s correlation coefficients, is insufficient. Additional tests such as the Box-Cox lambda transformation series, and synergy with axomyelin unit biology, provide more compelling evidence of the axon-fiber diameter relation.

A summary of Box-Cox transformations of the axon diameters from mouse A in the Sham EAE cohort (Fig. 1) is plotted against cognate fiber diameters in Fig. S1A. The color scheme of the data correspond with the lambda (λ) values of the *y*-axis transformations in decreasing order and the linear regression fits are extrapolated to intersect the *x*- and *y*-axes. Summary graphs of the transformed data from mice B and C (not shown) are equivalent.

The next step is generating and comparing the Spearman’s and Pearson’s correlation coefficients, *r_S_* and *r_xy_*, respectively, for each curve (see Methods). The Spearman’s coefficients of the Box-Cox transformations for mice A-C are: mouse A, *r_S_* = 0.988; mouse B, *r_S_* = 0.985; mouse C, *r_S_* = 0.986. The Pearson’s coefficients ±95% CIs of all transformations for the cohort are plotted as a function of λ in Fig. S1B, and only transformations with λ = 0.5 and λ = 1 conform to the Spearman’s versus Pearson’s criterion, *r_xy_* > *r_S_*.

In Fig. S1B, the lowest values of *r_xy_* and the widest CIs (the least likely to be correct), are from the λ = −3 transformation (inverse cube). The values of *r_xy_* increase and crest at λ = 1, decreasing thereafter. Accordingly, the *r_xy_* maximum corresponds with the original axon-fiber diameter correlation plot [i.e. (*y*-axis)^1^] and suggests that the axon-fiber diameter relation, as measured by the Pearson’s correlation coefficient (with the underlying assumption of normal distributions), is most closely satisfied with the linear regression fit in Fig. 2A. Moreover, a mixed-effects analysis using Fisher’s Z scores of the Pearson’s coefficients shows that the original linear regression fit (λ = 1) is statistically superior to all nonlinear fits from post hoc Dunnett’s tests, (0.001 < P < 0.028, λ = {-3, −2, −1, −0.5, 0, 0.5, 2}).

In addition to demonstrating that a linear relation for the axon-fiber diameter plot is statistically superior to a nonlinear relation, several biological properties also add to the constraints. Thus, when considering only myelinated axons, fiber diameters are larger by definition. Accordingly, extrapolations of the regression fits must intersect the *x*-axis at a point where D_F_ ≥ 0, and the *y*-axis at D_A_ ≤ 0. The *x*- and *y*-intercepts ±95% CI from Fig. S1A are plotted against the cognate lambda values (Fig. S1C), and the combination of these constraints excludes all transformations for −3 < λ < 0.5. In addition, the regression lines plotted must have slope < 1 (Fig. S1D), which excludes values of λ > 1. Together, the Pearson’s coefficient statistics and the biological constraints exclude all regression fits except for λ = 1, the linear regression fit, thereby demonstrating the axon-fiber diameter relation is directly proportional.

## Acknowledgements

This work was supported by a grants Boost Award from Wayne State University. I thank the organizers of the 53^rd^ American Society for Neurochemistry annual meeting, Lexington, Kentucky, USA, March 18-22, 2023 in the colloquium entitled “History, Theory, Applications, Use, and Misuse of the *g*-ratio”, which began this wonderful journey. I thank my colleagues Dr Anne Boullerne, Dr Doug L Feinstein and Dr Jeff L Dupree for access to raw data from their 2015 EAE study, insightful discussions, support and comradery. All aspects of this manuscript were designed, performed, interpreted and written by AG. AG declares there are no conflicts of interest.

---

**Figure S1.**
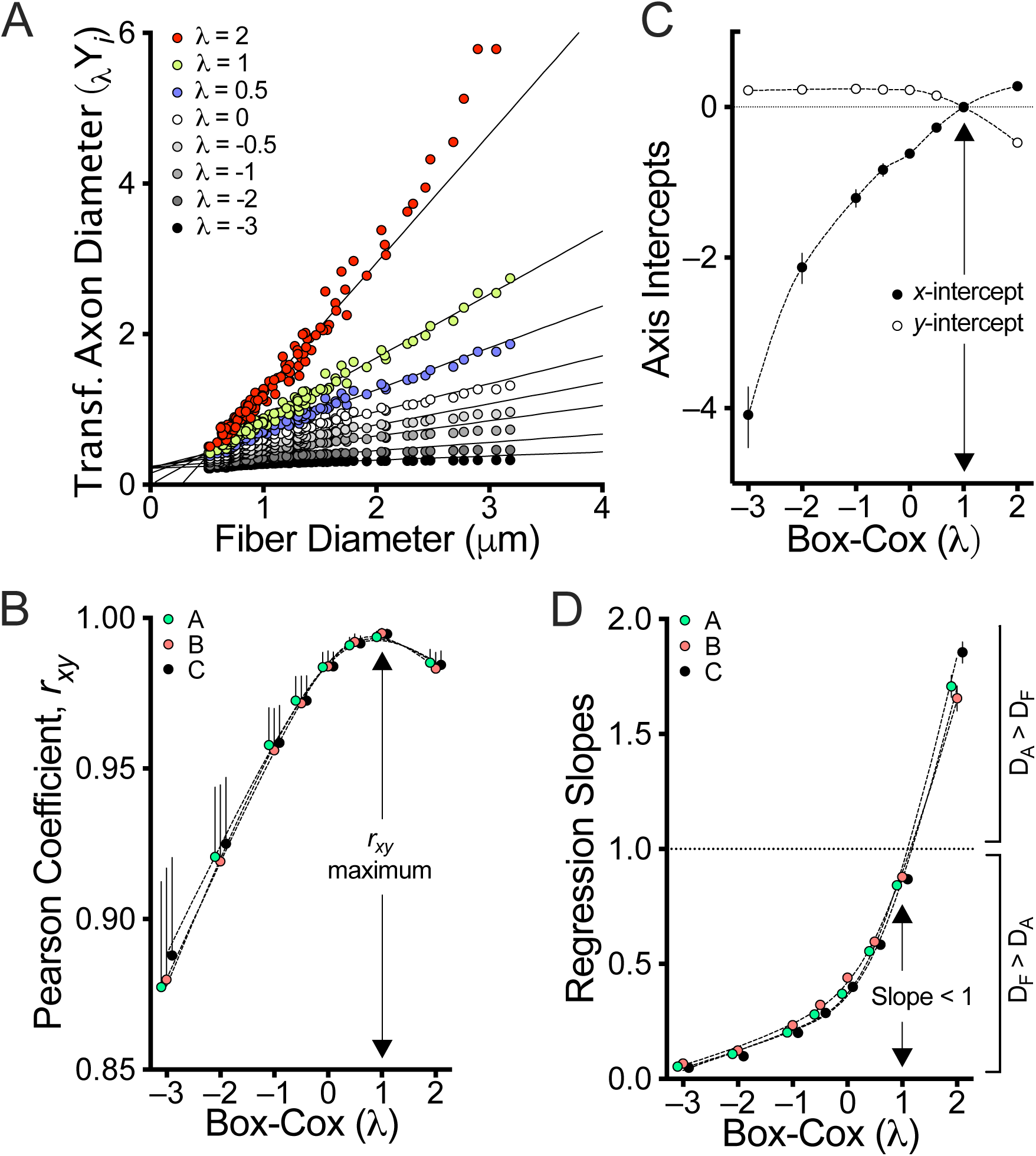
Statistical and biological constraints on axon-fiber diameter plots demonstrates a direct proportionality relation for the Sham (EAE) cohort **A.** Summary of a Box-Cox lambda transformation series for axon diameters from mouse A plotted against cognate fiber diameters (Fig. 1). The different transformations are fit using linear regression, and extrapolated to the *x*- and *y*-axes to estimate several parameters that identify the most likely axon-fiber diameter relation. **B.** Pearson’s correlation coefficients for the regression fits (mice A-C, Fig. 1C) are plotted against the corresponding Box-Cox lambda values. The maximum *r_xy_* value, for λ = 1, is the most likely axon-fiber diameter relation as confirmed by mixed-effects analysis with Geisser-Greenhouse correction (F_(1.23,2.65)_ = 1010, ε = .18, P = .0003). Post hoc Sidak’s multiple comparison’s tests show that no other values of lambda yield superior correlations. **C.** The *x*- and *y*-axis intercepts ±95% CI for the extrapolated regression fits in (A) plotted against the Box-Cox lambda values. Biological constraints on these intercepts exclude regression fits for all *x*-intercepts ≤ 0 and *y*-intercepts > 0, i.e. for all transformations of D_A_ where λ < 1. **D.** The constraints of myelinated fibers mandate that the regression slope < 1 (i.e. D_F_ > D_A_), which excludes regression fits where λ > 1.

